# Deep Learning for genomic prediction accounting for heterosis in crossbreeding systems

**DOI:** 10.1101/2025.07.24.666530

**Authors:** F. Shokor, P. Croiseau, H. Gangloff, S. Fritz, A.A.A. Martin, T. Mary-Huard, B.C.D. Cuyabano

## Abstract

**Background:** Crossbreeding is used in animal breeding to combine desirable traits from different breeds and to exploit hybrid vigor, and many approaches have been proposed to improve the prediction of genetic values of crossbred animals. This study aimed to assess whether deep learning methods can enhance the prediction of genetic values in crossbred populations by effectively capturing heterosis. We tested several prediction models that varied in how they incorporate genetic information, including purebred and crossbred data, breed composition, and breed-of-origin of alleles, on both simulated and real crossbred data. Finally, we also proposed an approach in which additive genetic effects and heterosis are predicted separately and then combined using effect-specific weights.

**Results:** Our results show that the presence of heterosis reduced the accuracy of traditional statistical methods but improved the performance of deep learning models. When heterosis was present, the highest prediction accuracy for total genetic value was achieved by combining additive genetic values, predicted using statistical method, with the predicted heterosis values, using a weighted sum. The explicit prediction of heterosis using deep learning and a regression method yielded similar results. The integration of breed-of-origin of alleles achieved the highest prediction accuracy for the additive genetic effect, outperforming both methods based on breed composition, and those that train the GBLUP model by combining purebred and crossbred animals. When applied to real data, adding the predicted heterosis did not increase prediction accuracies of crossbred animals, a result that we attributed to the homogeneity of the real crossbred population and its limited sample size.

**Conclusions:** The usefulness of deep learning for crossbred prediction depends on the degree of heterosis exhibited in the traits. Incorporating both additive genetic effects and heterosis considering the breed-of-origin of the alleles in the genomic information and effect-specific weighting leads to more accurate and robust predictions in our simulated data, particularly when crossbred data is heterogeneous. Even when prediction accuracy is not significantly improved, deep learning can still provide valuable insight into the degree of heterosis across traits, offering a deeper understanding of crossbred genetic architecture.

## Background

Improving the productivity, health, and sustainability of livestock and crop populations remains a central goal in breeding programs. As global demands increase and breeding challenges become more complex, innovative strategies are required to maximize genetic progress while preserving diversity and long-term viability (Sørensen et al., 2008; Wu & Zhao, 2021). Among these strategies, crossbreeding has gained significant attention in recent years and has been widely adopted across many species in animal and plant breeding programs (Berry, 2021; Labroo et al., 2021; VanRaden et al., 2020; Wu & Zhao, 2021). This growing interest in crossbreeding is primarily driven by the benefits of breed complementarity, i.e. the ability to combine desirable traits from distinct breeds, and the potential to retain heterosis (Wakchaure et al., 2015). Heterosis, or hybrid vigor, traditionally refers to a superior performance due to heterozygous genotypes at causal loci. However, in the context of crossbreeding, non-additive effects can also arise when alleles inherited from different breeds interact due to differences in their breed-of-origin and genetic background, even if the genotype at a locus appears homozygous. This is because the surrounding haplotypes and breed-specific linkage disequilibrium patterns may differ, influencing gene expression and trait outcomes. This effect enhances the performance of subsequent generations of the crossbred animals, with potential to significantly influence traits with low heritability such as fertility, health, and survival (Hansen, 2006).

In France, there is a growing interest in applying crossbreeding and rotational crossbreeding, a system where different breeds are used in succession over generations to maintain heterosis, in dairy cattle, involving the main dairy breeds, but particularly the French Holstein, Normande, and Montbéliarde. Although crossbreeding appears to be a promising approach to improve the genetic value of populations, its application remains limited. One reason for this limitation is the lack of evaluation methods for crossbred animals, necessary to retain these animals in breeding programs, and keep them in rotational crossbreeding systems. Therefore, currently the main focus remains on first-order crosses (F1), which are mostly used as terminal crosses and tend to capture the greatest benefits of heterosis. Nonetheless, the growing interest in a continuous crossbreeding system in France, and other countries, has driven the recent development of tools that would allow crossbred animals to be evaluated and selected more effectively.

A critical matter for selecting crossbred animals is the ability to accurately predict the genetic merit and phenotypic performance of crossbred individuals. In this context, genomic prediction (GP), which uses dense molecular markers to estimate genetic value (Meuwissen et al., 2001), initially raised high expectations for improving prediction across breeds. Genomic data offered the promise of capturing genetic differences directly at the genome level, enabling more precise evaluations. However, early attempts to predict crossbred or across-breed performance using purebred-derived genomic information often yielded limited success. Biological and statistical challenges, including differences in linkage disequilibrium (LD) structure, quantitative trait loci (QTL) effects, and allele frequencies, undermined the transferability of marker effects across breeds (de Roos et al., 2008; Olson et al., 2012; Steyn et al., 2019). Furthermore, given that genetic correlations for the same trait measured between purebred and crossbred populations are generally less than one, relying solely on purebred data to predict crossbred performance is often inadequate, especially when correlations are low (Barani et al., 2024; Dekkers, 2007).

To address these limitations in crossbreeding prediction, researchers have explored multiple strategies to enhance prediction accuracy of such animals, by optimizing both the composition of the reference population and the modeling approach. Including crossbred individuals alongside purebreds in the training set allows prediction models to better reflect the genetic structure of the target population. This approach has consistently increased prediction accuracy, particularly when the reference crossbred animals are closely related to those being evaluated (Barani et al., 2024; Bonifazi & Aivazidou, 2024; Cesarani et al., 2024; Esfandyari et al., 2015; Karaman et al., 2021; Londoño-Gil et al., 2025; Ma et al., 2024; Winkelman et al., 2015). On the modeling side, one approach is to treat animals from different breeds as distinct traits in a multi-breed evaluation framework (Bonifazi & Aivazidou, 2024; Olson et al., 2012; van den Berg et al., 2020; Vitezica et al., 2016). A more straightforward strategy to account for breed differences involves including global breed composition in the prediction model. This involves estimating the proportion of the genome inherited from each parental breed and using it to inform predictions (VanRaden et al., 2020).

While global breed composition provides a useful summary of genetic background, it does not capture the local ancestry of alleles, i.e., the breed of origin for each specific genomic segment. To overcome this limitation, breed-of-origin of alleles can be incorporated into prediction models. This method involves phasing crossbred genotypes and comparing them to purebred references to trace each allele back to its breed source. Rather than estimating SNP effects directly in crossbred populations, the model applies SNP effects previously estimated in purebreds, assigning them based on the breed-of-origin of each allele. This allows the model to account for breed-specific genetic architectures, including distinct LD patterns and allele frequencies. Incorporating such breed-aware information has been shown to improve prediction accuracy, especially when parental breeds are genetically divergent (Eiríksson et al., 2021; Guillenea et al., 2023; Rio et al., 2020; Saintilan et al., 2022).

Beyond breed structure, another layer of complexity in crossbred prediction arises from non-additive genetic effects, such as dominance and epistasis. While additive effects are often sufficient for selecting crossbred individuals as breeding candidates, non-additive effects play a key role in explaining phenotypic performance, particularly in the presence of heterosis. Including non-additive effects can improve prediction accuracy by capturing additional sources of genetic variation and enabling a more accurate partitioning of the total genetic variance into additive and non-additive components. Several studies have demonstrated that explicitly accounting for non-additive effects enhances crossbred prediction accuracy (González-Diéguez et al., 2021; Kristensen et al., 2023; Roth et al., 2022; Vitezica et al., 2016; Xiang et al., 2016; Zeng et al., 2013; Zhuo et al., 2024).

Capturing such complex genetic relationships has motivated increasing interest in machine learning (ML) and deep learning (DL) approaches for genomic prediction. These methods are well-suited for learning nonlinear, high-order interactions in large genomic datasets without relying on strict assumptions about underlying genetic models (Alzubaidi et al., 2021; Sarker, 2021). In particular, deep learning, which uses multi-layer neural networks to extract hierarchical features from raw input data, has shown potential for capturing both additive and non-additive effects (Abdollahi-Arpanahi et al., 2020; Bellot et al., 2018; Chen et al., 2024; Jubair et al., 2021; Lee et al., 2023; Shokor et al., 2025; K. Wang et al., 2023; Zingaretti et al., 2020). Although still in its early stages in crossbreeding applications, some promising results have emerged. For instance, Tusell et al., (2020) used a support vector machine (SVM)—a type of ML model—to predict crossbred pig performance and demonstrated that incorporating purebred genotype and phenotype data can enhance prediction accuracy for crossbred individuals.

The objectives of this study are to evaluate whether DL methods can improve the prediction of crossbred performance by better modeling heterosis effects. Specifically, we investigated whether DL can outperform traditional models by capturing heterosis effects under different levels, and whether it provides practical advantages when applied to real-world crossbred populations.

## Methods

To evaluate genomic prediction strategies in crossbred populations, we simulated a structured population derived from two genetically distinct purebred lines. These two breeds (Breed 1 and Breed 2) were subject to selection over 25 generations, each for a different trait to reflect divergent breeding objectives. After this purebred phase, a crossbreeding scheme was introduced, forming a crossbred population through rotational mating between the two breeds for an additional 10 generations. This simulation setup includes both additive and heterosis effects and serves as the foundation for evaluating multiple prediction models.

We simulated four quantitative traits with varying heritabilities and genetic correlations to represent different genetic architectures. For crossbred individuals, we explicitly modelled heterosis effects with different levels of contribution to total genetic variance.

### Simulated data

Each purebred population was initially composed of 1,000 individuals, resulting in 2,000 animals in the base generation. Data was simulated using package GenEval R (https://github.com/bcuyabano/GenEval, Cuyabano, 2021). 10,000 single nucleotide polymorphisms (SNPs) distributed across 29 chromosomes were simulated in LD (de Roos et al., 2008). To reflect genetic divergence between breeds, each purebred population was initialized with different allele frequencies for each SNP.

For each trait, the phenotypic data of purebred samples were simulated as

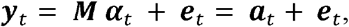

where ***M*** is a matrix of which element *M*_*i,j*_ corresponds to the centered genotype of individual *i* at QTL *j*, ***α***_t_ =[*α*_*t*,1_….*α*_*t*,q_] is the vector of QTL additive effects for trait *t =* 1,…,*T;****a***_*t*_ is the vector of true additive genetic values for trait t; and ***e***_*t*_ is the vector of residual errors. A total of q=500 QTLs were simulated, and the same set of QTL positions was used for all traits and both breeds.

The additive QTL effects *α*_*b,t,j*_ were initially sampled from a multivariate normal Distribution *α*_*b,t,j*_ ∼ *N*(0,ρ_*QTL*_⊗ ρ_*g*_), where ρ_*QTL*_ is a 2 × 2 breed-level QTL correlation matrix (diagonal = 1, off-diagonal = 0.3), and ρ_*g*_ is a 4 × 4 genetic correlation matrix defining the desired correlations between traits: 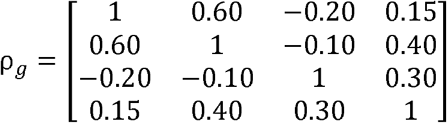.The Kronecker product structure results in an 8 × 8 covariance matrix, and ensures that QTL effects differ across traits and breeds, while maintaining the specified correlation structure. These genetic correlations were included to reflect biologically realistic relationships between traits, such as antagonistic and synergistic effects often observed in livestock breeding. While not explicitly modeled in the prediction frameworks, they contribute to the genetic architecture and affect selection responses across generations.

After sampling, the additive QTL effects were standardized and rescaled so that the resulting additive genetic variance for trait *t* equal to 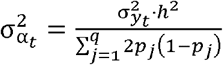, such that *h*^2^ is the narrow-sense heritability, 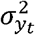 is the phenotypic variance, and *p*_*j*_ is the allele frequency 0 at QTL *j* in the initial (generation 0) purebred population. Only additive genetic effects were considered in purebred populations. The vector of the true genetic values (TGV) was computed as ***a***_*t*_ = ***M*α****_*t*_. Given the assumptions of the model, the resulting genetic values ***a***_*t*_ follow a multivariate normal distribution with 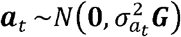 and 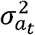 is the additive genetic variance; and 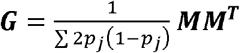 is the genomic relationship matrix; 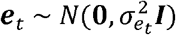 is the vector of random errors where 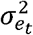 is the error variance. The traits were simulated each with different narrow-sense heritability (*h*^2^) (Trait 1: 0.3, Trait 2: 0.05, Trait 3: 0.1, and Trait 4: 0.5). For the base population, generation 0, all trait means were fixed at 0, and the phenotypic variance was standardized to 1 across all breeds and traits.

The two purebred populations were independently evolved for 25 generations, with selection applied only to males based on their phenotypic performance. Selection for Breed 1 was based on Trait 1 (*h*^2^= 0.3), while selection for Breed 2 was based on Trait 3 (*h*^2^= 0.1), with 50% of the top males retained in Breed 1 and 65% retained in Breed 2. No selection was applied to females. Selected males were used for four generations, while females were used for two generations. At each generation, 1,000 new individuals were generated per breed, maintaining a population structure of 50% females and 50% males.

A crossbred population was introduced in generation 25, allocating 10% of the top-selected males and females from each purebred population to crossbreeding; females selected for crossbreeding were no longer available for purebred mating. The crossbred population was then evolved for 10 additional generations, following a structured mating scheme. At each generation, the top 50% of crossbred females were selected based on their overall performance across all four traits, calculated as an unweighted average (equal weight of 1 for each trait). These selected females were then mated with purebred males from one pure breed per generation, producing second-order crosses. As a result, each generation contained a mix of new purebred individuals (females and males) from both breeds, new first-order crossbred individuals from both purebred, and new second-order crossbred individuals from one purebred. The breed of origin of each allele was tracked for crossbred individuals.

The phenotype for crossbred individual *i* for trait *t* was modeled as ***Y***_*t,i*_ = ***G***_*t,i*_ + ***e***_*t,i*_ = ***a***_*t,i*_ + ***h***_*t,i*_ + ***e***_*t,i*_, where ***G***_*t,i*_ is the vector of total genetic value. The additive genetic value for trait *t* in crossbred individual *i* was computed as: 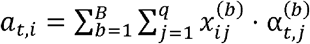, where **a**_t,i_ is the additive genetic value of individual *i* for trait *t, B* is the number of breeds, *q* is the number of QTL, 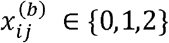 is the number of alternative alleles from breed *b* at QTL *j* in individual *i*,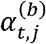 is the additive effect at QTL *j* for breed *b*, and the breed *b* is determined for each allele at each locus via breed-of-origin tracking.***e***_*t,i*_ represents the error vector simulated the same way as in purebred. The term ***h***_*t,i*_ represents the vector of heterosis effects introduced into the total genetic value for crossbred individual *i*. A heterosis effect was considered present when the QTL was heterozygous, and the two alleles originated from different breeds (e.g., A0B1 or A1B0). Both configurations were treated symmetrically and assigned the same heterosis effect, regardless of the breed of origin of the reference or alternate allele. While this assumption simplifies the modeling of heterosis, it does not account for potential allele-specific dominance or directional effects between breeds, where the effect of carrying an allele may differ depending on the breed it originates from, which may occur in real biological systems.

The heterosis effects dd of QTLs were initially sampled from an exponential distribution with *λ* = 10, producing small and strictly positive values. To ensure that heterosis effects remained within a biologically reasonable range, the values were scaled so that the maximum heterosis effect did not exceed the absolute value of the maximum additive effect (│*α*│).In addition, 30% of QTLs were assigned a dominance value of zero, so that only a subset of loci contributed to heterosis. The total heterosis effect (*h*) was computed as 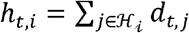 where ℋ_*i*_ is the set of QTL loci in individual *i* that are heterozygous and whose two alleles originate from different breeds, *d*_*t,j*_ is the heterosis effect at QTL *j* for trait *t*.

Although the individual heterosis effects were small and sparsely distributed, their variance was further scaled to achieve a predefined heterosis ratio *η* between heterosis variance and total genetic variance, defined as *η* = *Var*(*h*)/(*Var*(*a*)+ *Var*(*h*)). This allows precise manipulation of the relative contribution of heterosis to the total genetic variance at each generation. We chose to fix this ratio across generations to test a predefined and controlled heterosis-to-additive contribution, even though in real-life scenarios, this ratio may vary due to selection. However, accounting for such variation was beyond the scope of this study.Different heterosis ratios (*η*) were tested: *η=0* (no heterosis effect), *η* = 0.1 (low heterosis effect), *η* = 0.2 (moderate heterosis effect), and *η* = 0.3 (high heterosis effect). To assess the repeatability of our findings, 20 replicates of the complete dataset were simulated.

### Real Data

For the real data analysis, we used crossbred data from the French national database by selecting animals with both genotype and phenotype records whose ancestry involved the three main French dairy breeds: Holstein, Montbéliarde, and Normande. This resulted in a dataset of 1,358 F1 crossbred individuals, including 315 Montbéliarde × Holstein and 1,043 Normande × Holstein crosses.

To determine the breed of origin of each allele in the crossbred individuals, we used the BreedOrigin program (Saintilan et al., 2022). This method assigns breed origin to haplotypes, in a probabilistic manner, using sliding windows of n markers (16, 8, 4, 2, and 1), based on haplotype frequencies in reference purebred populations. Haplotype libraries were built using 5,000 individuals per breed: 2,000 selected based on the number of offspring (to capture widely represented haplotypes), and 3,000 randomly chosen among animals born between 2023 and 2025 (to capture recent genetic variation). Additionally, 230 Red Holstein bulls were included to represent the genetic diversity of this subpopulation. Breed-of-origin assignment was enforced at every marker. When the probabilistic method could not confidently determine the origin of a segment, its breed was imputed based on the origin of surrounding segments.

Phenotypic data included first-lactation records for four traits: milk yield, fat yield, protein yield, and somatic cell score (SCS). SNP effects for these traits in the three pure breeds were obtained from breed-specific single-step evaluations. We used the mean of the traits for each breed as reported in Table 1 of Dezetter et al. (2015).

**Table 1:** Data requirements for different prediction models for crossbred populations.

| Model Name | Genotype<br>CB | Genotype<br>PB | SNP Effects PB | BOA<br>Data | Breed<br>Proportion | Modeling<br>Heterosis |
| --- | --- | --- | --- | --- | --- | --- |
| GBLUP CB | Yes |  |  |  |  |  |
| GBLUP ALL | Yes | For training |  |  |  |  |
| DL CB | Yes |  |  |  |  | Implicitly |
| DL ALL | Yes | For training |  |  |  | Implicitly |
| Global Admixture | Yes |  | Breed-specific | From pedigree |  |  |
| Local Admixture Pure | Yes | For BOA | Breed-specific | Yes |  |  |
| Local Admixture joint | Yes | For BOA | Joint breed | Yes |  |  |
| Heterosis Regression | Optional | Optional |  | Optional | From pedigree | Explicitly |
| DL Heterosis | Yes | For BOA |  |  | Yes | Explicitly |
| Weighted Sum: Local Admixture + Regression Heterosis | Yes | For BOA | Breed-specific | Yes |  | Explicitly |
| Weighted Sum: Local Admixture + DL Heterosis | Yes | For BOA | Breed-specific | Yes |  | Explicitly |

### Statistical Methods for genomic prediction

To predict the genetic values of crossbred individuals, we compared two main strategies. In the first approach, we predicted crossbred genetic values by training a Genomic Best Linear Unbiased Prediction (GBLUP) model using available genotype - without QTLs - and phenotype data. The GBLUP model was applied in two reference population (training set): (i) using only crossbred individuals, and (ii) using all individuals jointly, including both purebred and crossbred animals. The model is defined as:

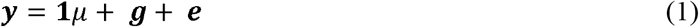

where ***y*** is the phenotype vector; **1** is a vector of ones; *μ* is the overall mean; ***g*** is the vector of genetic values, with 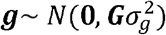: ***G*** is the genomic relationship matrix (GRM) (VanRaden, 2008) and 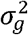 is the genetic variance. The vector of random errors ***e*** is assumed to follow 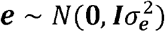, in which 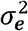 is the error variance and ***I*** is the identity matrix.

In the second approach, we predicted the genetic values of crossbred individuals using their admixture structure and SNP effects estimated from purebred populations. Two different strategies were used to estimate the genetic value of crossbred:

1. Global admixture model:
  - This approach follows the method proposed by VanRaden et al. (2020), which uses the genomic breed composition - referred to as Breed Base Representation (BBR)- of crossbreds, following the model:

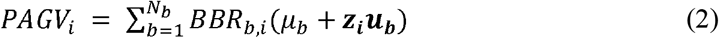
  - where *PAGV*_*i*_ is the predicted additive genetic value of animal *i*, BBR_*b,i*_ is the BBR–breed composition- value for breed *b* in animal *i*, ***z***_***i***_is a vector of allele contents for animal *i*; *μ*_*b*_ is the overall mean for breed *b*. ; and ***μ***_***b***_ is the vector of estimated SNP effects for breed *b*. These ***μ***_***b***_ were estimated with breed-specific analysis, in which GBLUP models—following Equation (1)—were applied independently to each purebred population. The SNP effects ***μ***_*b*_ were computed by back-solving from ***g***_*b*_ as described in H. Wang et al. (2012).
2. Local admixture model:
  - To account for local ancestry, we implemented the model proposed by Eiríksson et al. (2021), which uses phased genotypes and breed-of-origin (BOA) information at each locus. The predicted additive genetic value for individual ii is calculated as:

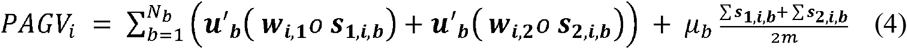

where *N*_*b*_ is the number of breeds. The term ***w***_*i,j*_ is the vector of phased data for haplotype *j* (where *j* = 1 or 2) of animal *i*, with alleles coded as 0 or 1 for alternative alleles; ***s***_***j***,***i***,***b***,_ is the breed indicator vector indicating the origin of alleles in haplotype *j* of animal *i* where a value of 1 is assigned to alleles from breed *b*, and 0 indicates an allele from the alternative other breed. The ***μ***_***b***_ vector can be obtained either from the breed-specific analysis (resulting in predicted values denoted *PAGV_i_pure_*), or from a joint-breed analysis (denoted *PAGV*_*i_joint*_). The joint model, based on Karaman et al. (2021), is defined as:

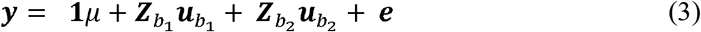

where ***y*** represents the phenotype vector for all individuals and all breeds; **1** is a vector of 1s, and *μ* is the general mean. The vectors 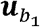 and 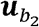 correspond to the estimated SNP effects for breed 1 and 2 respectively, while 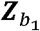 and 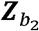 are the matrices containing breed-specific SNP content for breed 1 and 2, respectively, coded as described in Guillenea et al. (2023).

We used the R (4.1.1) package BGLR package (1.0.8) (Pérez & de los Campos, 2014) to fit the statistical models on the simulated data. We ran the models with 30,000 iterations, discarding the first 10,000 samples as burn-in with a thinning 10.

### Heterosis regression

The heterosis coefficient (H) was estimated using breed-of-origin of alleles (BOA) information. For each individual, we calculated the proportion of SNPs for which the two alleles originated from different breeds. This direct method captures the actual genomic heterozygosity due to crossbreeding and does not rely on parental breed proportions.

As an alternative, Dechow et al. (2007) proposed estimating *H* from the global breed proportions (admixture) of the sire and dam: 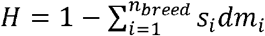, where *s*_*i*_ and *dm*_*i*_ are the proportions of sire and dam SNP from breed *i*, respectively.

After computing *H*, the mean heterosis can be estimated by regressing on *H* following the approach of Dezetter et al. (2015). The individual heterosis effect for each animal is then calculated by multiplying its heterosis coefficient *H* by the estimated mean heterosis effect.

### Deep Learning Model

We proposed two deep learning (DL) models to predict the genetic performance of crossbred individuals, differing in how they account for heterosis. The first model is a classical Multi-Layer Perceptron (MLP) that implicitly captures heterosis effects through its nonlinear modeling capacity. The second model is designed to account for heterosis explicitly, by encoding genomic features that reflect breed-of-origin and heterozygosity at the SNP level.

For the first model, we investigated the effect of training set composition by comparing two reference population structures: one using only crossbred individuals (DL CB) and another combining purebred and crossbred individuals (DL ALL). To predict trait values in crossbred animals, we trained a Multi-Layer Perceptron (MLP) on each of these training sets (Goodfellow et al., 2016). The input to the network was a vector *x* ∈ *R*^*m*^, where *m* corresponds to the number of SNP markers. Each SNP genotype was encoded as 0, 1, or 2, corresponding to the number of alternate alleles (i.e., homozygous reference = 0, heterozygous = 1, homozygous alternate = 2). This numeric encoding allows the model to learn additive effects across the genome.

In the second DL model, we focused on predicting the heterosis genetic value for each crossbred individual. The key idea was to explore alternative ways of encoding the genotype data so that the input representation explicitly captures the genomic signals responsible for heterosis, namely, heterozygosity and breed-of-origin of alleles. Unlike the first model, which implicitly captured both additive and non-additive effects, this model was designed to predict heterosis effects explicitly. The resulting predicted heterosis values were later combined with predicted additive values from the first model to generate overall trait predictions.

To predict the heterosis genetic value we coded the genomic data of crossbred samples using different encodings:

1. BOA Encoding: Each SNP marker is coded as 1 if the two alleles at the locus originate from different breeds, and 0 if both alleles originate from the same breed. This encoding captures breed divergence due to crossbreeding but does not capture the actual allelic state or zygosity.
2. Dominance Encoding: In this approach, SNPs are coded based solely on allelic state: heterozygous genotypes are coded as 1, and homozygous genotypes are coded as 0. This encoding captures the classical dominance deviation resulting from heterozygosity, independent of breed origin.
3. Combined BOA-Dominance Encoding: This encoding integrates both heterozygosity and breed-of-origin information. A SNP is coded as 1 only if it is heterozygous and the two alleles originate from different breeds; otherwise, it is coded as 0. This encoding corresponds directly to the heterosis mechanism implemented in the simulation, where a heterosis effect is introduced at a QTL only when both heterozygosity and inter-breed origin of alleles are present. Intra-breed heterozygosity does not contribute to heterosis, and the direction of allele inheritance (e.g., Breed 1 vs. Breed 2 origin) is not distinguished in this encoding.
4. One hot Encoding: The genomic data of each crossbred individual was encoded into a matrix of dimension *m* × 5. The five encoded features represent distinct allele configurations: the first feature encodes SNPs where both alleles originate from the same breed (always coded as 0), while the remaining four represent different configurations in which alleles originate from different breeds, both alleles being 0, both alleles being 1, and cases where the allele ‘1’ comes from either breed 1 or breed 2. This encoding method provides a detailed representation of heterosis effects by distinguishing between homozygous and heterozygous allele combinations across breeds and identifying the breed of origin of reference alleles. This enables the model to learn distinct genetic effects associated with breed-specific allele inheritance patterns.

Each encoding strategy was evaluated for its effectiveness in predicting heterosis genetic values and capturing the non-additive genetic effects that contribute to crossbred performance. The tested encoding strategies were designed to capture inter-breed heterosis, that is, heterosis arising from allele combinations inherited from different breeds. These encodings do not account for potential heterosis effects within the same breed (intra-breed heterozygosity) in crossbred individuals.

Detailed information on the network architecture, activation functions, and layer sizes is provided in Appendix 1.

### Estimation of Trait-Specific Weights for Additive and Heterosis Effects

To optimally combine additive and heterosis genetic value predictions, we estimated trait-specific coefficients using a linear regression approach. For each trait, the final phenotype *y* was modeled as a weighted sum of the predicted additive (*PAGV*) and heterosis (*PHGV*) genetic values:

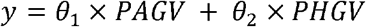

where *θ*_1_ and *θ*_2_ are the estimated weights. These coefficients were obtained by solving the normal equations of least squares:

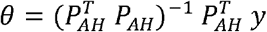

where *P*_*AH*_ represents the matrix stacking additive and heterosis predictions across individuals,and 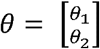 the learned weights.

This two-step approach, first estimating additive and heterosis values independently, then combining them, is the core strategy used to generate final predictions in our framework. It provides a flexible and data-driven way to capture the relative contribution of additive and non-additive genetic components for each trait.

Table 1 summarizes the data required by each tested model to predict the genetic value of crossbred individuals.

### Model Training

All the DL models were trained by minimizing the Huber loss (Hastie et al., 2001) across the four traits, for 100 maximum epochs. Each epoch consisted of several gradient updates performed on mini-batches of size 200, covering the entire training dataset once per epoch. An early stopping rule was employed to terminate training if there was no improvement in validation loss after 10 epochs. Adam optimizer was used for the training with a learning rate of 10^−4^ and then reduced to 10^−5^ once the loss plateaued. The choice of optimizer, learning rate, activation function and number of neurons in hidden layers was made after a grid search. The model was implemented using Pytorch (1.10.2) in Python (3.11.6) and trained on a single GPU with 48 GB memory (NVIDIA A40).

### Data Split

To train and evaluate the deep learning (DL) models, the dataset was split into three sets based on generations: a training set (PB: generations 21 to 27, 6,300 samples per purebred; CB: generations 25 to 27, 1,400 samples), a validation set (generations 28 and 29, 1,200 samples), and a test set (generations 30 and 31, 1,200 samples). The test set corresponds to individuals for whom predicted genetic values (PGV) are to be obtained in the absence of phenotypic records, while both the training and validation sets consist of individuals with available genotypes and phenotypes. Unlike statistical models, DL requires an internal validation set with complete information to fine-tune model parameters and prevent overfitting. In addition to hyperparameter tuning, the validation set was also used to estimate the weights for combining additive and heterosis effects, instead of using the training set, since it is the closest in terms of generation to the test set. For the GBLUP model, the training and validation sets were merged into a single training set to fit the model, and the same test set was used for evaluation.

In the real dataset, animals born in the last 2 years two years (2016–2017; 224 samples) were used for test. The older samples formed the training set, and the most recent individuals (born in 2015; 368 samples) from the training set (766 samples) were set aside as the validation set for the DL models.

### Evaluation metric

To evaluate model performance, we used the Pearson correlation between the predicted genetic values and the true simulated total genetic values as a measure of prediction accuracy. For the heterosis model, the correlation was calculated between the predicted and true heterosis values when the heterosis ratio was different from zero. For real data, the prediction accuracy was calculated as the Pearson correlation between the phenotype and the predicted genetic value.

## RESULTS

In this section, we present and compare the results of the simulation and the different models described earlier. The reported results are based on the test set, which includes generations 30 and 31.

### Simulation results

Figure 1 shows the genetic progress of purebred and crossbred animals over generations. Progress is measured as the difference between each generation’s mean genetic value and that of generation 0. Traits under selection improved more in their respective breeds: Trait 1 in Breed 1 and Trait 3 in Breed 2. Due to their negative correlation (–0.20), gains in one led to declines in the other. Trait 2, positively correlated with Trait 1 (0.60), showed higher gains in Breed 1. Trait 4, not directly selected but positively correlated with Traits 1 (0.15) and 3 (0.30), improved similarly in both breeds.

**Figure 1.**
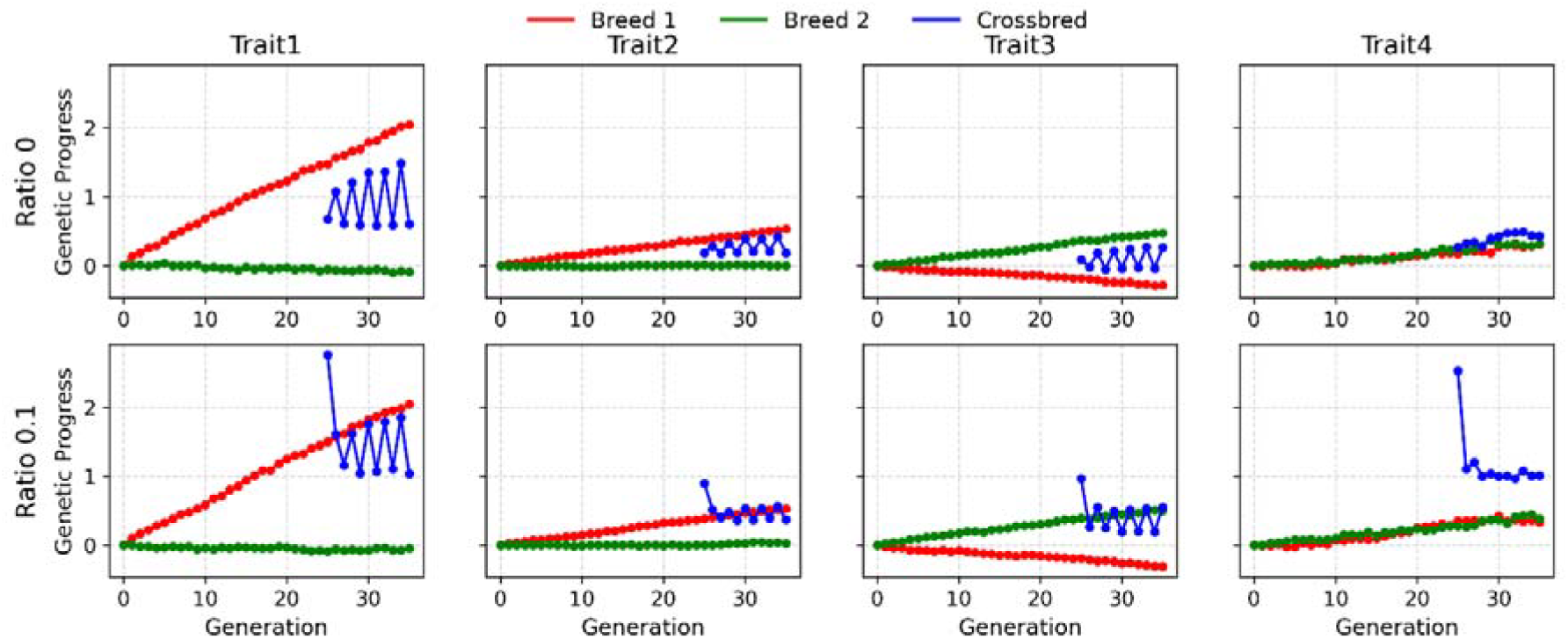
Genetic progress over generations for four traits in two purebred lines (Breed 1 in red, Breed 2 in green) and their crossbred progeny (in blue), under two heterosis ratio scenarios (0 and 0.1). Each subplot shows the genetic gain per generation for Trait 1 to Trait 4. The top row represents a heterosis ratio (variance heterosis / variance total genetic value) of 0, while the bottom row corresponds to a ratio of 0.1.

In crossbreds, females were selected on all traits. Without heterosis, their genetic values were intermediate. With heterosis, they improved and approached the superior purebred, peaking in the first crossbred generation.

Crossbred progress varied by which breed was used for mating. Generation 26, sired by Breed 1 males, had higher Trait 1 values than Generation 27, sired by Breed 2. The reverse was seen for Trait 3. For Trait 4, crossbreds outperformed both breeds due to heterosis and balanced parental contributions. The Principal Component Analysis (PCA) supported this pattern: purebreds were clearly separated, and crossbreds shifted toward the sire’s breed across generations, reflecting backcrossing and breed composition changes (see Appendix 2).

### Crossbred Prediction Performance

Figure 2.A represents the prediction accuracy of total genetic value (ToGV) from GBLUP and DL models trained on simulated data using only crossbred (CB) and all breeds (ALL) across different heterosis ratios (0, 0.1, 0.2, and 0.3). Since the results on simulated data were computed using the true simulated genetic values rather than phenotypes, we expected higher accuracy values. These values are not directly constrained by trait heritability and can therefore range between 0 and 1, unlike results based on phenotypes, which are limited by the proportion of genetic variance explained. In general, the prediction accuracy of ToGV from GBLUP models decreased as the heterosis ratio increased, however the prediction accuracy of ToGV from DL models increased as the heterosis ratio increased. Across all scenarios, models trained on data from all breeds consistently outperformed those trained on crossbreds only. At a heterosis ratio of 0, GBLUP ALL achieved the highest accuracy across all traits (0.81, 0.62, 0.66, 0.70). As the heterosis ratio increased, deep learning models—particularly DL ALL—showed improved performance and began to outperform the GBLUP models, with gains ranging from 0.06 to 0.25 at a heterosis ratio of 0.3. At the same time, the performance gap between GBLUP ALL and GBLUP CB narrowed, with the average difference across traits decreasing from 0.15 at ratio 0 to 0.09 at ratio 0.3. At higher heterosis levels, DL ALL consistently delivered the highest prediction accuracy across all traits (at ratio 0.3: 0.75, 0.51, 0.65, 0.71), followed by GBLUP ALL (at ratio 0.3: 0.63, 0.45, 0.51, 0.46). The DL CB model showed the lowest prediction accuracy at heterosis ratio 0 but gradually improved as the heterosis ratio increased, eventually reaching a performance level similar to the GBLUP models at heterosis ratio 0.3.

**Figure 2.**
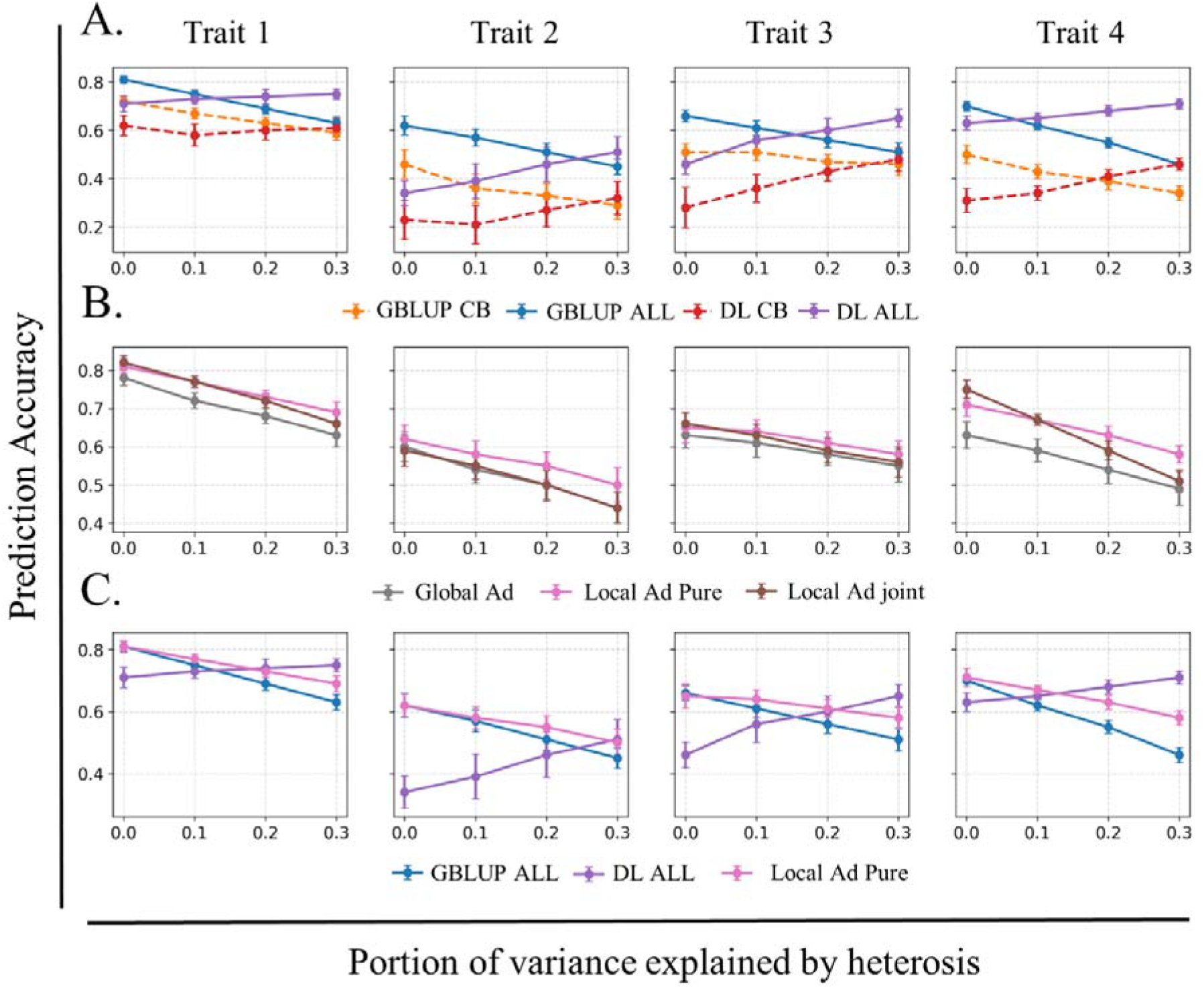
Prediction accuracy of total genetic value from models: A. GBLUP CB (trained with crossbred only), GBLUP ALL (trained with crossbred and purebred), DL CB, and DL ALL; B. Global Ad(mixture), Local Ad(mixture) Pure (breed specific SNP effects), and Local Ad(mixture) joint (joint SNP effects); C. GBLUP ALL, DL ALL, and Local Ad Pure, across heterosis ratios (0, 0.1, 0.2, 0.3). Each subplot represents a trait, showing model performance trends as heterosis ratio varies. A. Solid lines represent models trained on all data (purebred and crossbred), while dashed lines represent models trained on crossbred data only. The dots represent the mean of prediction accuracy calculated across 20 replicas and error bars indicate the standard deviation

By comparing the models that account for global (breed proportion) and local (breed-of-origin of alleles (BOA)) admixture (Figure 2.B), all three approaches exhibited a progressive decline in prediction accuracy of ToGV as the heterosis ratio increased. The global admixture model consistently yielded the lowest prediction accuracy across all traits and heterosis ratios. Models incorporating BOA demonstrated a similar performance across all scenarios. At higher heterosis ratios, the local admixture model using SNP effects from breed-specific (pure) models (at ratio 0.3: 0.69, 0.50, 0.58, 0.58) showed a slight advantage over the joint evaluation model (joint) (at ratio 0.3: 0.66, 0.44, 0.56, 0.51). When compared to GBLUP and DL models trained with all breeds, this model showed a higher prediction accuracy than GBLUP model when heterosis ratio increased and similar to DL model except when the heterosis ratio was too high (0.3) (Figure 2.C).

Overall, DL model showed higher prediction accuracy than statistical methods when the heterosis ratio increased. However, for Trait 2 (h^2^ = 0.05), DL model yielded performance similar to Local Admixture Pure model.

### Heterosis prediction

Figure 3 compares the prediction accuracy of heterosis value (HV), measured as the correlation between the true heterosis genetic value and the predicted value using DL model, from different SNP encoding strategies: 1-hot, BOA, DOM, and BOA× DOM, and the regression heterosis method, across all traits and heterosis ratios (0.1, 0.2, and 0.3). Overall, the regression method performed similarly to the best DL model using BOA encoding, and even outperformed it in the scenario with low heritability (Trait 2) and low heterosis. Among the DL encoding strategies, the BOA encoding consistently outperformed the other methods across all scenarios. The 1-hot and BOA × DOM encodings showed similar performance. The DOM encoding performed the worst under all conditions, showing the lowest accuracy overall. However, incorporating dominance effects into BOA (BOA × DOM) led to better accuracy than using DOM alone, but lower accuracy than using BOA alone. The prediction accuracy of HV from all models improved as both the heterosis ratio and heritability increased.

**Figure 3.**
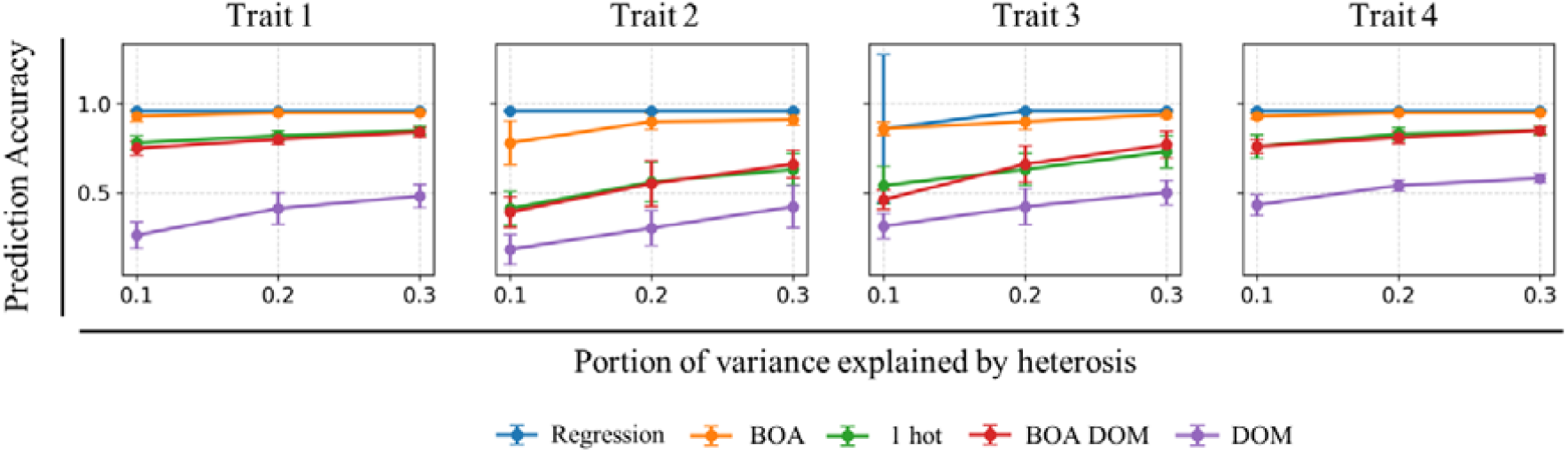
Prediction accuracy of heterosis genetic value from DL models using different SNP encoding methods—1-hot, BOA, DOM, and BOA+DOM— and the heterosis regression method across heterosis ratios (0.1, 0.2, and 0.3). BOA refers to breed-of-origin allele encoding, DOM encodes dominance effects, and 1 hot refers to the detailed encoding of homozygosity and heterozygosity of crossbred.

### Weighted sum

Figure 4 presents a comparison between five models: the additive model (Local Admixture with breed-specific effects), the DL heterosis model using BOA encoding, the unweighted sum of additive and DL heterosis predictions, the weighted sum of the additive predictions and the DL heterosis predictions and the regression heterosis prediction. Results are shown across different heterosis ratios. For each scenario, prediction accuracy is reported as the correlation between the predicted and true values for total genetic effect.

**Figure 4.**
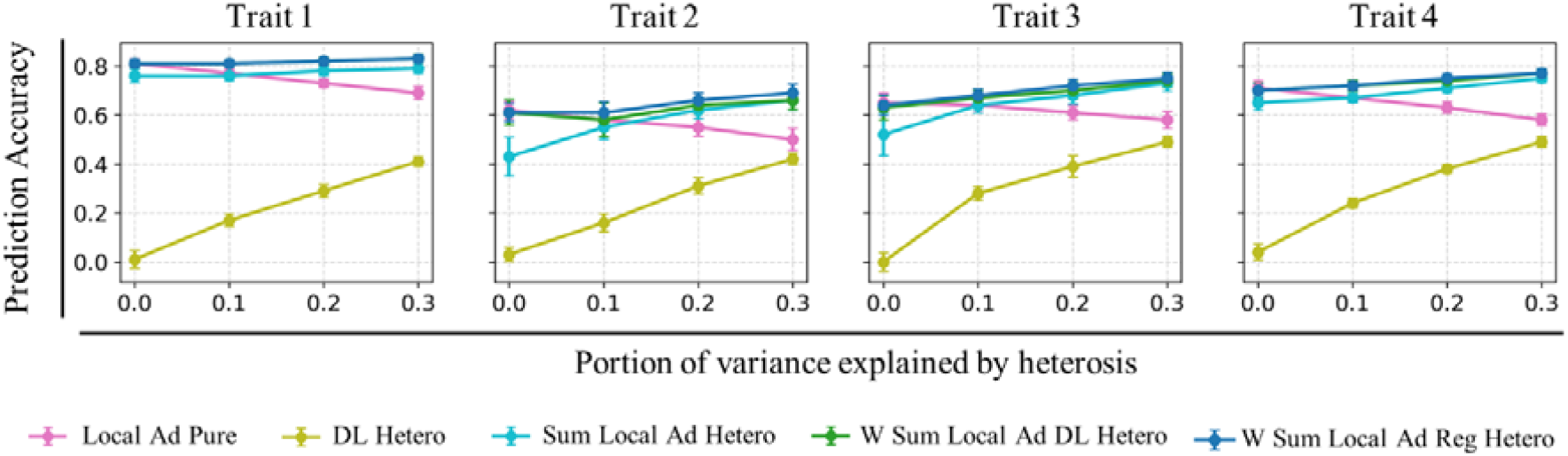
Prediction accuracy of total genetic values from model: Local Ad(mixture) pure (Breed Specific), Heterosis with BOA encoding, sum Local Ad Pure and Heterosis, Weighted Sum Local Ad Pure and DL Heterosis, and Weighted Sum Local Ad Pure and regression Heterosis. Results are shown across different heterosis ratios (0, 0.1, 0.2, and 0.3).

Adding the predicted heterosis value, from DL or regression method, to the predicted additive value in a weighted sum consistently resulted in the highest prediction accuracy of the total genetic value. When the true genetic effect was purely additive, incorporating the predicted heterosis value without applying weights reduced the accuracy of the total genetic value. However, when estimated weights were used, the prediction accuracy of the ToGV did not decrease, compared to the additive model alone, although no improvement was observed either.

At low heterosis ratio, both the additive model and the weighted sum model yielded similar prediction accuracies of ToGV. Nonetheless, the weighted sum model provided a better estimation of the heterosis effect than the additive model. As the heterosis ratio increased, the prediction accuracy of the additive model for the ToGV began to decline.

In contrast, the weighted sum model maintained stable the total prediction accuracy across all heterosis ratios. This stability was achieved by a trade-off: as the heterosis ratio increased, the additive effect decreased while the heterosis effects increased. The unweighted sum model (i.e., simple addition without weights) also showed an improved total accuracy with increasing heterosis ratio, eventually reaching a similar level to the weighted sum model. However, it consistently exhibited higher heterosis representation.

The heterosis-only model produced the lowest accuracy for both total and additive genetic values, while achieving the highest representation of the heterosis effects.

### Real Data

On real data, we applied the methods that performed best on simulated data: Local Ad(mixture) Pure and the weighted sum approach using heterosis predictions from both the DL and regression methods. The results showed similar performance across all methods. However, adding the heterosis value in a weighted sum to the additive predictions did not improve the prediction accuracy of the genetic value (Table 2).

**Table 2:** Prediction accuracy of genetic value from model: Local Ad(mixture) pure (Breed Specific), Weighted Sum Local Ad Pure and DL Heterosis, and Weighted Sum Local Ad Pure and regression Heterosis.

|  | Local Ad(mixture)<br>Pure | Weighted Sum Local Ad Pure<br>and regression Heterosis | Weighted Sum Local Ad Pure and<br>DL Heterosis |
| --- | --- | --- | --- |
| **Milk Yield** | 0.42 | 0.41 | 0.41 |
| **Fat Yield** | 0.37 | 0.36 | 0.36 |
| **Protein Yield** | 0.42 | 0.41 | 0.41 |
| **SCS** | 0.30 | 0.30 | 0.30 |

## Discussion

In this study, we evaluated whether deep learning (DL) methods could improve the prediction of crossbred performance by effectively modeling heterosis effects. Using simulated data, we compared the performance of various statistical methods that utilize different sources of information, purebred and crossbred genotypes, breed proportions (global admixture), and breed-of-origin of alleles (BOA) (local admixture), to predict the genetic value of crossbred individuals. We also evaluated the performance of a DL model that accounts for heterosis either implicitly or explicitly through the use of a heterosis encoding scheme; in the explicit case, the model can estimate an individual specific heterosis genetic value. Our results showed that the presence of heterosis reduced the prediction accuracy of all tested statistical methods, while it improved the accuracy of the DL models. Additionally, we proposed an approach to combine the additive predicted genetic value from statistical method with the heterosis value predicted from DL, or from regression through a weighted sum. This proposed method achieved the highest prediction accuracy for total genetic value in scenarios where heterosis was present.

Since crossbred populations are often smaller in size, compared to purebreds, relying solely on crossbred data to predict genetic values can result in a poor estimation of allelic effect. One strategy to enhance prediction accuracy is to incorporate data from the purebred populations from which the crossbred animals are derived. In line with previous studies (Barani et al., 2024; Misztal et al., 2022), we demonstrated that this approach improved the prediction accuracy of crossbred individuals across all scenarios, including varying heritabilities and levels of heterosis (Figure 2). However, when heterosis effects were strong, the advantage of the GBLUP model trained on combined purebred and crossbred data over the model trained on crossbreds alone diminished. This suggests that the reduced benefit may result from the inability of GBLUP, which models primarily additive effects, to capture the increasing contribution of non-additive (heterosis) effects in these scenarios.

The statistical model that incorporated breed-of-origin of alleles (BOA) showed higher prediction accuracy than the GBLUP model and models based on breed proportion (Figure 2). The BOA model also demonstrated less sensitivity to heterosis effects compared to other statistical approaches, with an average drop in prediction accuracy of 0.11 between traits with a heterosis ratio of 0 and 0.3, compared to a drop of 0.19 observed with GBLUP All. While GBLUP models and global admixture-based approaches can capture general patterns of relatedness between purebred and crossbred populations, local admixture-based models offer a more precise representation of the breed origin of each allele. This finer resolution allows local admixture models to better capture heterosis effects when present, resulting in higher prediction accuracy under such conditions, showing an average improvement of 0.08 across traits compared to GBLUP ALL. However, when no heterosis was simulated, the performance of the local admixture model was similar to that of GBLUP ALL, indicating that its main advantage lies in scenarios where non-additive effects contribute significantly to genetic value.

However, how SNP effects are estimated in purebred populations plays a crucial role. Guillenea et al. (2023) found that joint analysis outperforms breed-specific evaluations, a pattern that we also observed in our results when heterosis is absent. Conversely, in the presence of heterosis, especially for traits with low heritability (e.g., Trait 2), SNP effects estimated from breed-specific evaluations appear to be more effective. Joint evaluations estimate SNP effects by allowing interactions between breeds but implicitly assume a large additive genetic architecture. This may limit their ability to capture non-additive effects. In contrast, breed-specific evaluations preserve the ancestral context of allele effects without enforcing interdependences across breeds. Since crossbred individuals often exhibit non-additive effects distinct from both parental lines, SNP effects estimated independently for each breed may better reflect the deviations observed in crossbreds relative to the purebred mean. This could explain why the local admixture model using breed-specific (pure) SNP effects outperformed the model using joint-breed SNP effects (Figure 2.B).

It is well known that the prediction accuracy of genomic models is positively related to the heritability of the trait of interest (Frouin et al., 2020). Higher heritability implies a larger proportion of phenotypic variance attributable to genetic variance, which typically leads to higher prediction accuracy, given a population size compared to traits with lower h^2^. This relationship is reflected in our results: Trait 1 (h^2^ = 0.3) showed the highest prediction accuracy, followed by Trait 3 (h^2^ = 0.1), with the lowest accuracy observed for Trait 2 (h^2^ = 0.05). However, heritability is not the only factor influencing prediction accuracy. When comparing Trait 1 (h^2^ = 0.3) and Trait 4 (h^2^ = 0.5), we observed that Trait 4 had lower prediction accuracy despite its higher heritability. This highlights the influence of selection history on prediction performance.

In a rotational crossbreeding scheme where each generation of crossbred animals is back-crossed to a different purebred line, the genetic mean of crossbred animals for the selected trait tends to change in favor of the contributing purebred. Alternating the breed used in each generation introduces fluctuations in the genetic mean across crossbred generations— not due to heritability, but due to opposing selection pressures in the two parental lines. Prediction accuracy is influenced by the variance of genetic values within the population. When selection has been applied differently across purebred populations, this leads to divergence in allele frequencies, creating an additional source of genetic variance in the crossbred population. As a result, prediction accuracy tends to be higher for traits under divergent selection. In contrast, traits that were not directly selected in either purebred population, but are positively correlated with selected traits (such as Trait 4), tend to show similar genetic trends across both purebreds. This limits the divergence and, consequently, reduces the genetic variance available for prediction in the crossbreds. Consequently, crossbred individuals show minimal divergence in genetic values for that trait (trait 4), and the observed genetic variance, and thus prediction accuracy, is mostly dictated by heritability alone. Finally, although Trait 2 was also not directly selected for in either purebred, it was positively correlated with Trait 1 (0.6) and negatively correlated with Trait 3 (-0.1). These correlations introduced indirect selection effects, generating some variation across purebreds and crossbreds and thereby influencing prediction accuracy beyond what heritability alone would suggest.

Deep learning (DL) is well known for its ability to detect non-linear patterns in data, which makes it particularly suited for capturing heterosis effects in traits. This was confirmed in our study, where the DL model, trained solely on genomic data without explicitly encoding heterosis, outperformed the GBLUP model in scenarios with substantial heterosis, with gains in prediction accuracy reaching up to 0.25 (Trait 4 with ratio 0.3). These results highlight the potential of DL to extract non-additive signals directly from genomic features. However, when the goal is to estimate heterosis effects more precisely, for example, to interpret their biological or economic relevance, it is preferable to use input data that explicitly encodes heterosis, such as local breed-of-origin information. In our experiments, incorporating such information further improved both the accuracy and interpretability of heterosis predictions.

Various definitions and encoding strategies for heterosis have been explored in our study. Since heterosis arises in crossbred animals due to differences in the breed of origin of alleles, encoding SNPs based on their breed-of-origin (BOA) enables the model to capture admixture effects, that emerge simply because the two alleles originated from different breeds. However, adding dominance information to the BOA encoding led to reduced prediction accuracy, even though heterosis in the simulated data was introduced by adding a heterosis effect when a QTL was heterozygous with alleles originating from different breeds. This discrepancy may be explained by the absence of actual QTLs in our input data. Even if a QTL is in linkage disequilibrium with nearby SNPs, heterozygosity at the QTL does not guarantee heterozygosity at the linked SNPs, leading to inconsistent representations and potentially reduced prediction accuracy. We also tested a one-hot encoding approach, which provides a more granular representation of heterosis—such as the specific effects of having reference alleles from one breed versus another. However, since our simulated data did not include such detailed breed-specific effects, the model using one-hot encoding underperformed compared to the general BOA encoding. This encoding method may prove itself useful in simulations where breed-specific allele effects are present. Nonetheless, one significant limitation of this approach is its dimensionality: the input size increases substantially with the number of purebred breeds, potentially complicating model training and interpretation.

To obtain a more representative estimate of the total genetic value that included both the additive and heterosis effects we proposed a weighted sum of predictions from the best-performing statistical method and the heterosis prediction. This weighted approach reflects the full genetic potential of crossbred individuals by adapting the contribution of each component. Unlike a simple unweighted sum, our approach adjusts the relative influence of additive and heterosis effects, using a linear regression trained on crossbred performance. This adaptive scheme maintained high prediction accuracy of the total genetic value across all traits and heterosis levels, including cases where heterosis was absent, as the model learned to assign small or no weight to the heterosis term when it was not informative. The resulting weights are interpretable, easy to estimate, and require no prior knowledge of the true heterosis level, making the approach robust and efficient. Moreover, the predicted heterosis values generated by the deep learning model can be combined with additive predictions from any statistical or machine learning method, allowing flexible integration into existing genomic prediction pipelines. This makes the framework broadly applicable for improving crossbred evaluations across a range of breeding programs.

When applied to the real data available to us, our approach did not show any improvement when adding the predicted heterosis genetic value to the additive PGV. Several factors may explain these results. First, the data we used contained only first-order crosses, meaning the individuals have similar admixture levels. As a result, the predicted heterosis values—being conditioned on admixture level—are also similar across all samples. Adding a similar value to all individuals does not impact correlation levels (Appendix 3), which highlights the importance of including crosses with diverse admixture levels in the dataset. Second, another limiting factor of our real dataset was its size. It is well known that prediction accuracy generally decreases with smaller datasets. While this affects most prediction models, DL methods are more sensitive to limited dataset sizes, even if it is diverse (Appendix 4).

In practice, the available real data suffers from limited phenotypic and genotypic records, with only first-order (F1) crosses. Our results on simulated data suggest that integrating second-order crosses is crucial for improving prediction accuracy, by capturing heterosis effects, as long as they are well represented. Otherwise, including heterosis values for samples with small variation will not affect overall predictions and will not influence selection decisions.

The deep learning model contributed to a better understanding of the role of heterosis in crossbred performance. By leveraging regression-based techniques, DL provided valuable insights into the genetic architecture of crossbred individuals by quantifying the contribution of heterosis across traits. While the deep learning model did not consistently improve prediction accuracy over regression-based methods in this case, simulation results showed that under strong heterosis scenarios, the DL-ALL model can outperform traditional methods, even without explicitly modeling heterosis.

Finally, models that rely on previously estimated SNP effects from purebreds, rather than large genotype datasets for each purebred population are becoming more practical in terms of data efficiency and computational cost. In our study, we used only 5,000 genotyped purebred individuals to infer the breed-of-origin of alleles (BOA) in crossbred animals. In contrast, estimating breed-specific SNP effects required a much larger dataset of over 100,000 purebred animals. This illustrates that BOA inference can be achieved with a relatively small subset of data, making such models more accessible in scenarios where purebred genotyping is limited.

In this study, we applied methods that consider the breed of origin of alleles and require the presence of genotyped crossbred animals. However, since larger datasets often include ungenotyped crossbreds, alternative approaches that incorporate such samples may be advantageous. One such method is the ssGBLUP with Metafounders (Legarra et al., 2015), in which pseudo-individuals are assigned to the base population. This method accounts for both phenotyped and genotyped samples, while also modeling the correlation between subpopulations and the metafounders. This concept has shown promising results when applied to crossbred populations, especially when dominance effects are included (Poulsen et al., 2022; Zhuo et al., 2024). However, when complete data are available, that is, when both purebred parents and their crossbred offspring are genotyped and phenotyped, the advantages of this method compared to alternative genomic prediction models become limited. For this reason, it was not considered in our analysis. Nonetheless, in real-world scenarios where missing data are common and purebreds may be related, applying this method could improve prediction accuracy (Barani et al., 2024).

## Conclusions

In this study, we tested many approaches to predict the genetic value of crossbred individuals in the presence of heterosis effects. Using breed-of-origin of alleles (BOA) information, a deep learning (DL) model was proposed to predict sample-specific heterosis values, achieving performance comparable to a regression-based heterosis method. These heterosis predictions were then combined, via a weighted sum, with additive genetic values predicted by a statistical method. This approach consistently yielded the highest prediction accuracy across all tested scenarios. We demonstrated the importance of incorporating both purebred and crossbred data in the evaluation process. Furthermore, our results showed that using BOA information to estimate genetic values led to more accurate predictions. We also observed that the presence of heterosis reduced the prediction accuracy of statistical methods (GBLUP-based), while enhancing the performance of deep learning models. Accounting for heterosis improved the prediction of total genetic value in crossbred animals, highlighting its potential to enhance selection strategies. This improvement became more pronounced as the heterogeneity of the crossbreeding dataset increased.

## Declarations

### Ethics approval and consent to participate

Not applicable

### Consent for publication

Not applicable

### Availability of data and materials

All source code in this study are freely available at:

https://github.com/fshokor/DL_CrossB_Heterosis

## Authors’ contributions

FS designed the deep learning models and the data simulation method, performed all the analysis, and did the main writing of the manuscript. PC participated in the conceptualization of the study and the discussion of the results, and in revising the manuscript. HG participated in the design of the deep learning models, and in revising the manuscript. SB participated in the discussion of the results. AAAM prepared the real data and performed BOA analysis, and participated in revising the manuscript. TMH participated in designing the data simulation method, in the design of the deep learning models, and in revising the manuscript. BCDC participated in the conceptualization of the study, in designing the data simulation method, in the discussion and interpretation of the results, and in revising the manuscript. All authors read and approved the final manuscript.

## Acknowledgements

This work was supported by INRAE Metaprogramme DIGIT-BIO (Digital biology to explore and predict living organisms in their environment). We would like to think GenIALearn team for their valuable discussion and contribution, and Jocelyn de Goer De Herve from (INRAE /UMR EPIA) for managing the server on which DL work was done.

## Additional files

**Appendix 1**

The first model consisted of the following components:

Input layer: Receives a vector *x* ∈ *R*^*m*^, where *m* is the number of SNP markers. SNP genotypes were encoded as 0 (homozygous reference), 1 (heterozygous), or 2 (homozygous alternate), representing the number of alternate alleles per marker.

Hidden Layer 1: A fully connected layer with 400 neurons, followed by a LeakyReLU activation function with a negative slope of 0.1.

Hidden Layer 2: A fully connected layer with 256 neurons, again followed by a LeakyReLU activation.

Output Layer: A fully connected linear layer that outputs predicted genetic values for the four traits.

The forward pass of the network can be described as follows:

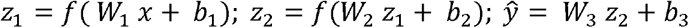

where *W*^*i*^ ∈ *R*^*h*×*m*^ and *b*_*i*_ ∈ *R*^*m*^ are the parameters, weights and biases, respectively, of the first and second hidden layer and the output layer with *i* = 1,2,and 3, respectively.

The second prediction model consists of multiple (four) hidden layers with LeakyReLU activations after each linear layer, followed by dropout regularization, which helps prevent overfitting and improves generalization across traits. The resulting intermediate representation is then passed through a linear layer that outputs a multi-trait prediction (predicted heterosis genetic value PHGV).

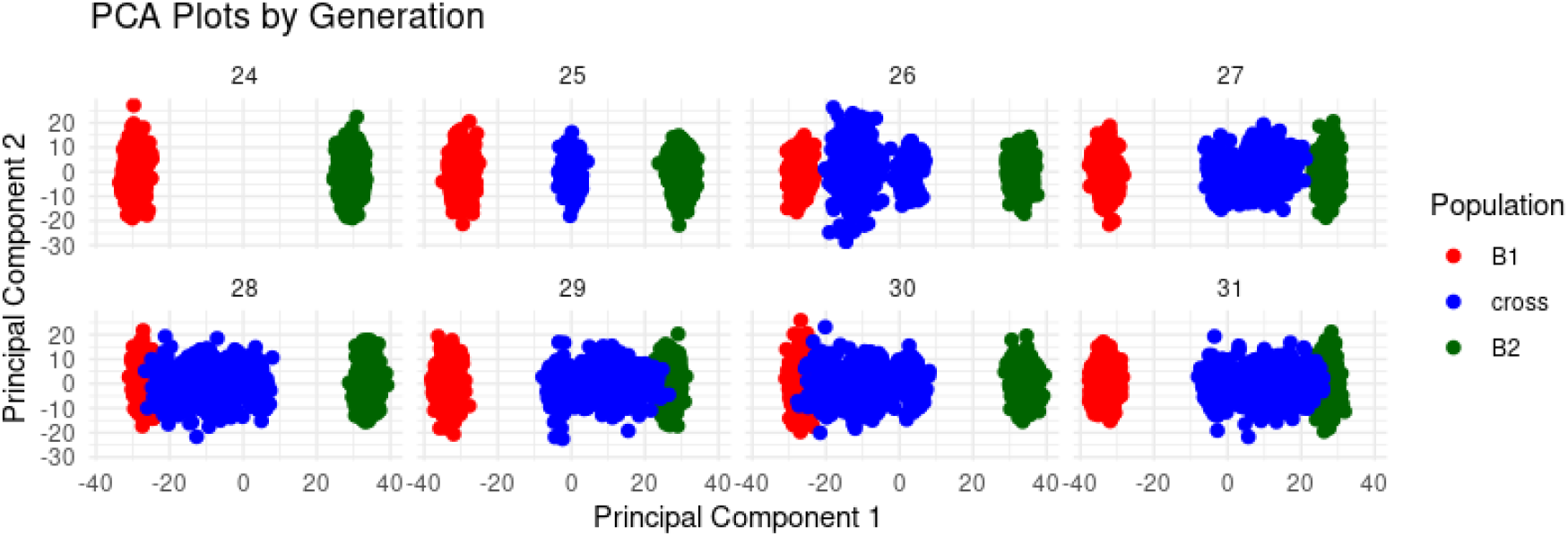

*Appendix 2: Principal Component Analysis (PCA) of genotype data from individuals of Breed 1 (red), Breed 2 (green), and their crossbred progeny (blue) across generations 24 to 31. The plot illustrates the genetic structure and differentiation among populations over time. Crossbreds occupy an intermediate position between the two purebreds, with shifts in distribution across generations reflecting alternating mating schemes and changes in breed composition.*

*Appendix 3: Table presenting the results of predictions made using simulated data with a total of 1,300 samples, for heterosis ratio of 0.1. It compares the prediction accuracy of genetic values from the following models: Local Ad(mixture) Pure (Breed-Specific), Weighted Sum of Local Ad Pure and DL-predicted Heterosis, and Weighted Sum of Local Ad Pure and Regression-predicted Heterosis. Prediction accuracy was calculated as the Pearson correlation between the predicted values and the phenotypes in the test set.*

|  | Local Ad(mixture)<br>Pure | Weighted Sum Local Ad Pure and<br>regression Heterosis | Weighted Sum Local Ad Pure and<br>DL Heterosis |
| --- | --- | --- | --- |
| **Trait 1** | 0.27 | 0.27 | 0.27 |
| **Trait 2** | 0.21 | 0.24 | 0.2 |
| **Trait 3** | 0.14 | 0.13 | 0.11 |
| **Trait 4** | 0.5 | 0.52 | 0.48 |

*Appendix 4: Table presenting the results of predictions made using simulated data, where samples were selected based on having more than 90% of SNPs from different breeds (i.e., high breed heterozygosity), for a heterosis ratio of 0.1. It compares the prediction accuracy of genetic values from the following models: Local Ad(mixture) Pure (Breed-Specific), Weighted Sum of Local Ad Pure and DL-predicted Heterosis, and Weighted Sum of Local Ad Pure and Regression-predicted Heterosis. Prediction accuracy was calculated as the Pearson correlation between the predicted values and the phenotypes in the test set.*

|  | **Local Ad(mixture)**<br>**Pure** | **Weighted Sum Local Ad Pure and**<br>**regression Heterosis** | **Weighted Sum Local Ad Pure and**<br>**DL Heterosis** |
| --- | --- | --- | --- |
| **Trait 1** | 0.32 | 0.32 | 0.32 |
| **Trait 2** | 0.11 | 0.1 | 0.11 |
| **Trait 3** | 0.12 | 0.12 | 0.12 |
| **Trait 4** | 0.5 | 0.5 | 0.5 |

## References

Abdollahi-Arpanahi, R., Gianola, D., & Peñagaricano, F. (2020). Deep learning versus parametric and ensemble methods for genomic prediction of complex phenotypes. Genetics Selection Evolution, 52(1), 12. 10.1186/s12711-020-00531-z

Alzubaidi, L., Zhang, J., Humaidi, A. J., Al-Dujaili, A., Duan, Y., Al-Shamma, O., Santamaría, J., Fadhel, M. A., Al-Amidie, M., & Farhan, L. (2021). Review of deep learning: Concepts, CNN architectures, challenges, applications, future directions. Journal of Big Data, 8(1), 53. 10.1186/s40537-021-00444-8

Barani, S., Miraie Ashtiani, S. R., Nejati Javaremi, A., Khansefid, M., & Esfandyari, H. (2024). Optimizing purebred selection to improve crossbred performance. Frontiers in Genetics, 15. 10.3389/fgene.2024.1384973

Bellot, P., de los Campos, G., & Pérez-Enciso, M. (2018). Can Deep Learning Improve Genomic Prediction of Complex Human Traits? Genetics, 210(3), 809–819. 10.1534/genetics.118.301298

Berry, D. P. (2021). Invited review: Beef-on-dairy—The generation of crossbred beef × dairy cattle. Journal of Dairy Science, 104(4), 3789–3819. 10.3168/jds.2020-19519

Bonifazi, R., & Aivazidou, S. (2024). Multi-breed multi-trait single-step genomic predictions for Holstein and Jersey including crossbred animals. 60.

Cesarani, A., Lourenco, D., Bermann, M., Nicolazzi, E. L., VanRaden, P. M., & Misztal, I. (2024). Singlestep genomic predictions for crossbred Holstein and Jersey cattle in the United States. JDS Communications, 5(2), 124–128. 10.3168/jdsc.2023-0399

Chen, C., Bhuiyan, S., Ross, E., Powell, O., Dinglasan, E., Wei, X., Atkin, F., Deomano, E., & Hayes, B. (2024). Genomic prediction for sugarcane diseases including hybrid Bayesian-machine learning approaches. Frontiers in Plant Science, 15. 10.3389/fpls.2024.1398903

de Roos, A. P. W., Hayes, B. J., Spelman, R. J., & Goddard, M. E. (2008). Linkage disequilibrium and persistence of phase in Holstein-Friesian, Jersey and Angus cattle. Genetics, 179(3), 1503– 1512. 10.1534/genetics.107.084301

Dechow, C. D., Rogers, G. W., Cooper, J. B., Phelps, M. I., & Mosholder, A. L. (2007). Milk, fat, protein, somatic cell score, and days open among Holstein, Brown Swiss, and their crosses. Journal of Dairy Science, 90(7), 3542–3549. 10.3168/jds.2006-889

Dekkers, J. C. M. (2007). Marker-assisted selection for commercial crossbred performance1. Journal of Animal Science, 85(9), 2104–2114. 10.2527/jas.2006-683

Dezetter, C., Leclerc, H., Mattalia, S., Barbat, A., Boichard, D., & Ducrocq, V. (2015). Inbreeding and crossbreeding parameters for production and fertility traits in Holstein, Montbéliarde, and Normande cows. Journal of Dairy Science, 98(7), 4904–4913. 10.3168/jds.2014-8386

Eiríksson, J. H., Karaman, E., Su, G., & Christensen, O. F. (2021). Breed of origin of alleles and genomic predictions for crossbred dairy cows. Genetics Selection Evolution, 53(1), 84. 10.1186/s12711-021-00678-3

Esfandyari, H., Sørensen, C., & Bijma, P. (2015). A crossbred reference population can improve the response to genomic selection for crossbred performance. Genetics Selection Evolution, 47. 10.1186/s12711-015-0155-z

Frouin, A., Dandine-Roulland, C., Pierre-Jean, M., Deleuze, J.-F., Ambroise, C., & Le Floch, E. (2020). Exploring the Link Between Additive Heritability and Prediction Accuracy From a Ridge Regression Perspective. Frontiers in Genetics, 11. 10.3389/fgene.2020.581594

González-Diéguez, D., Legarra, A., Charcosset, A., Moreau, L., Lehermeier, C., Teyssèdre, S., & Vitezica, Z. G. (2021). Genomic prediction of hybrid crops allows disentangling dominance and epistasis. Genetics, 218(1), iyab026. 10.1093/genetics/iyab026

Goodfellow, I., Bengio, Y., & Courville, A. (2016). Deep Learning. https://www.deeplearningbook.org/

Guillenea, A., Lund, M. S., Evans, R., Boerner, V., & Karaman, E. (2023). A breed-of-origin of alleles model that includes crossbred data improves predictive ability for crossbred animals in a multi-breed population. Genetics Selection Evolution, 55(1), 34. 10.1186/s12711-023-00806-1

Hansen, L. B. (2006). MONITORING THE WORLDWIDE GENETIC SUPPLY FOR DAIRY CATTLE WITH EMPHASIS ON MANAGING CROSSBREEDING AND INBREEDING.

Jubair, S., Tucker, J. R., Henderson, N., Hiebert, C. W., Badea, A., Domaratzki, M., & Fernando, W. G. D. (2021). GPTransformer: A Transformer-Based Deep Learning Method for Predicting Fusarium Related Traits in Barley. Frontiers in Plant Science, 12, 761402. 10.3389/fpls.2021.761402

Karaman, E., Su, G., Croue, I., & Lund, M. S. (2021). Genomic prediction using a reference population of multiple pure breeds and admixed individuals. Genetics Selection Evolution, 53(1), 46. 10.1186/s12711-021-00637-y

Kristensen, P. S., Sarup, P., Fé, D., Orabi, J., Snell, P., Ripa, L., Mohlfeld, M., Chu, T. T., Herrström, J., Jahoor, A., & Jensen, J. (2023). Prediction of additive, epistatic, and dominance effects using models accounting for incomplete inbreeding in parental lines of hybrid rye and sugar beet. Frontiers in Plant Science, 14. 10.3389/fpls.2023.1193433

Labroo, M. R., Studer, A. J., & Rutkoski, J. E. (2021). Heterosis and Hybrid Crop Breeding: A Multidisciplinary Review. Frontiers in Genetics, 12. 10.3389/fgene.2021.643761

Lee, H.-J., Lee, J. H., Gondro, C., Koh, Y. J., & Lee, S. H. (2023). deepGBLUP: Joint deep learning networks and GBLUP framework for accurate genomic prediction of complex traits in Korean native cattle. Genetics Selection Evolution, 55(1), 56. 10.1186/s12711-02300825-y

Legarra, A., Christensen, O. F., Vitezica, Z. G., Aguilar, I., & Misztal, I. (2015). Ancestral Relationships Using Metafounders: Finite Ancestral Populations and Across Population Relationships. Genetics, 200(2), 455–468. 10.1534/genetics.115.177014

Londoño-Gil, M., López-Correa, R., Aguilar, I., Magnabosco, C. U., Hidalgo, J., Bussiman, F., Baldi, F., & Lourenco, D. (2025). Strategies for genomic predictions of an indicine multi-breed population using single-step GBLUP. Journal of Animal Breeding and Genetics = Zeitschrift Fur Tierzuchtung Und Zuchtungsbiologie, 142(1), 43–56. 10.1111/jbg.12882

Ma, H., Li, H., Ge, F., Zhao, H., Zhu, B., Zhang, L., Gao, H., Xu, L., Li, J., & Wang, Z. (2024). Improving Genomic Predictions in Multi-Breed Cattle Populations: A Comparative Analysis of BayesR and GBLUP Models. Genes, 15(2), 253. 10.3390/genes15020253

Meuwissen, T. H. E., Hayes, B. J., & Goddard, M. E. (2001). Prediction of Total Genetic Value Using Genome-Wide Dense Marker Maps. Genetics, 157(4), 1819–1829. 10.1093/genetics/157.4.1819

Misztal, I., Steyn, Y., & Lourenco, D. A. L. (2022). Genomic evaluation with multibreed and crossbred data*. JDS Communications, 3(2), 156–159. 10.3168/jdsc.2021-0177

Olson, K. M., VanRaden, P. M., & Tooker, M. E. (2012). Multibreed genomic evaluations using purebred Holsteins, Jerseys, and Brown Swiss. Journal of Dairy Science, 95(9), 5378–5383. 10.3168/jds.2011-5006

Pérez, P., & de los Campos, G. (2014). Genome-Wide Regression and Prediction with the BGLR Statistical Package. Genetics, 198(2), 483–495. 10.1534/genetics.114.164442

Poulsen, B. G., Ostersen, T., Nielsen, B., & Christensen, O. F. (2022). Predictive performances of animal models using different multibreed relationship matrices in systems with rotational crossbreeding. Genetics Selection Evolution, 54(1), 25. 10.1186/s12711-022-00714-w

Rio, S., Moreau, L., Charcosset, A., & Mary-Huard, T. (2020). Accounting for Group-Specific Allele Effects and Admixture in Genomic Predictions: Theory and Experimental Evaluation in Maize. Genetics, 216(1), 27–41. 10.1534/genetics.120.303278

Roth, M., Beugnot, A., Mary-Huard, T., Moreau, L., Charcosset, A., & Fiévet, J. B. (2022). Improving genomic predictions with inbreeding and nonadditive effects in two admixed maize hybrid populations in single and multienvironment contexts. Genetics, 220(4), iyac018. 10.1093/genetics/iyac018

Saintilan, R., Croiseau, P., Baur, A., Croue, I., Ducrocq, V., Thomasen, J. R., Karaman, E., Boichard, D., Leclerc, H., & Cuyabano, B. C. D. (2022). Accuracy of prediction for a genomic evaluation in rotational crossbreeding scheme (Montbéliarde × Holstein × Red Danish). 6.

Sarker, I. H. (2021). Machine Learning: Algorithms, Real-World Applications and Research Directions. SN Computer Science, 2(3), 160. 10.1007/s42979-021-00592-x

Shokor, F., Croiseau, P., Gangloff, H., Saintilan, R., Tribout, T., Mary-Huard, T., & Cuyabano, B. C. D. (2025). Deep learning and genomic best linear unbiased prediction integration: An approach to identify potential nonlinear genetic relationships between traits. Journal of Dairy Science, 108(6), 6174–6189. 10.3168/jds.2024-26057

Sørensen, M. K., Norberg, E., Pedersen, J., & Christensen, L. G. (2008). Invited Review: Crossbreeding in Dairy Cattle: A Danish Perspective. Journal of Dairy Science, 91(11), 4116–4128. 10.3168/jds.2008-1273

Steyn, Y., Lourenco, D. A. L., & Misztal, I. (2019). Genomic predictions in purebreds with a multibreed genomic relationship matrix1. Journal of Animal Science, 97(11), 4418–4427. 10.1093/jas/skz296

Tusell, L., Bergsma, R., Gilbert, H., Gianola, D., & Piles, M. (2020). Machine Learning Prediction of Crossbred Pig Feed Efficiency and Growth Rate From Single Nucleotide Polymorphisms. Frontiers in Genetics, 11, 567818. 10.3389/fgene.2020.567818

van den Berg, I., MacLeod, I. M., Reich, C. M., Breen, E. J., & Pryce, J. E. (2020). Optimizing genomic prediction for Australian Red dairy cattle. Journal of Dairy Science, 103(7), 6276–6298. 10.3168/jds.2019-17914

VanRaden, P. M. (2008). Efficient Methods to Compute Genomic Predictions. Journal of Dairy Science, 91(11), 4414–4423. 10.3168/jds.2007-0980

VanRaden, P. M., Tooker, M. E., Chud, T. C. S., Norman, H. D., Megonigal, J. H., Haagen, I. W., & Wiggans, G. R. (2020). Genomic predictions for crossbred dairy cattle. Journal of Dairy Science, 103(2), 1620–1631. 10.3168/jds.2019-16634

Vitezica, Z. G., Varona, L., Elsen, J.-M., Misztal, I., Herring, W., & Legarra, A. (2016). Genomic BLUP including additive and dominant variation in purebreds and F1 crossbreds, with an application in pigs. Genetics Selection Evolution, 48(1), 6. 10.1186/s12711-016-0185-1

Wakchaure, R., Ganguly, S., Praveen, P., Sharma, S., Kumar, A., Mahajan, T., & Qadri, K. (2015). Importance of Heterosis in Animals: A Review. International Journal of Advanced Engineering Technology and Innovative Science, 1, 1–5.

Wang, H., Misztal, I., Aguilar, I., Legarra, A., & Muir, W. M. (2012). Genome-wide association mapping including phenotypes from relatives without genotypes. Genetics Research, 94(2), 73–83. 10.1017/S0016672312000274

Wang, K., Abid, M. A., Rasheed, A., Crossa, J., Hearne, S., & Li, H. (2023). DNNGP, a deep neural network-based method for genomic prediction using multi-omics data in plants. Molecular Plant, 16(1), 279–293. 10.1016/j.molp.2022.11.004

Winkelman, A. M., Johnson, D. L., & Harris, B. L. (2015). Application of genomic evaluation to dairy cattle in New Zealand. Journal of Dairy Science, 98(1), 659–675. 10.3168/jds.2014-8560

Wu, X.-L., & Zhao, S. (2021). Editorial: Advances in Genomics of Crossbred Farm Animals. Frontiers in Genetics, 12, 709483. 10.3389/fgene.2021.709483

Xiang, T., Christensen, O. F., Vitezica, Z. G., & Legarra, A. (2016). Genomic evaluation by including dominance effects and inbreeding depression for purebred and crossbred performance with an application in pigs. Genetics Selection Evolution, 48(1), 92. 10.1186/s12711-016-0271-4

Zeng, J., Toosi, A., Fernando, R. L., Dekkers, J. C., & Garrick, D. J. (2013). Genomic selection of purebred animals for crossbred performance in the presence of dominant gene action. Genetics Selection Evolution, 45(1), 11. 10.1186/1297-9686-45-11

Zhuo, Y., Du, H., Diao, C., Li, W., Zhou, L., Jiang, L., Jiang, J., & Liu, J. (2024). MAGE: Metafounders-assisted genomic estimation of breeding value, a novel additive-dominance single-step model in crossbreeding systems. Bioinformatics (Oxford, England), 40(2), btae044. 10.1093/bioinformatics/btae044

Zingaretti, L. M., Gezan, S. A., Ferrão, L. F. V., Osorio, L. F., Monfort, A., Muñoz, P. R., Whitaker, V. M., & Pérez-Enciso, M. (2020). Exploring Deep Learning for Complex Trait Genomic Prediction in Polyploid Outcrossing Species. Frontiers in Plant Science, 11, 25. 10.3389/fpls.2020.00025

